# Cell-free DNA Reveals Potential Zoonotic Reservoirs in Non-Human Primates

**DOI:** 10.1101/481093

**Authors:** Mark Kowarsky, Iwijn De Vlaminck, Jennifer Okamoto, Norma F Neff, Matthew LeBreton, Julius Nwobegabay, Ubald Tamoufe, Joseph Diffo Ledoux, Babila Tafon, John Kiyang, Karen Saylors, Nathan D Wolfe, Stephen R Quake

**Affiliations:** Department of Physics, Stanford University, Stanford, CA 94305.; Departments of Bioengineering and Applied Physics, Stanford University, Stanford, CA 94305.; Chan Zuckerberg Biohub, San Francisco, CA 94158.; Mosaic, BP 35353, Yaounde, Cameroon; Military Health Research Center (CRESAR), Carrefour Intendance, Yaounde, Cameroon; Metabiota Cameroon, Carrefour Intendance, Yaounde, Cameroon; Ape Action Africa, Yaounde, Cameroon; Limbe Wildelife Centre, Limbe, Cameroon; Metabiota, San Francisco, CA 94104; Global Viral, San Francisco, CA 94104.

## Abstract

The microbiome of non-human primates is relatively neglected compared with humans, and yet it is a source of many zoonotic diseases. We used high throughput sequencing of circulating cell-free DNA to identify the bacteria, archaea, eukaryotic parasites and viruses from over 200 individual non-human primates across 17 species from Africa. Many of the assembled sequences have low or no homology to previously sequenced microorganisms, while those that do have homology support prior observations of specific taxa present in primate microbiomes. The structure of the total microbiome is correlated with geographic location, even between distinct primate species which are co-located. However, we find that viruses have a particularly notable association with host taxa independent of geography. Numerous potentially zoonotic taxa were discovered in an unbiased manner, thereby expanding knowledge of host species diversity and strengthening the case for monitoring wildlife reservoirs.

**One Sentence Summary:** Blood from non-human primates provides insight into potential pathogens which might eventually infect humans.

## Main Text

Circulating nucleic acids in the blood contain molecules sampled from tissues throughout the body and have been used to monitor fetal development(*1*, *2*), organ transplants(*3* *4*), cancers(*5*) and infections(*6*, *7*). Our recent work has shown that *de novo* metagenomic assemblies allow the study of the microbiome in a hypothesis-free and unbiased manner, resulting in the discovery that the diversity of microbial life within humans is substantially larger than previously expected(*8*). This global approach augments the understanding of the human microbiome as determined from large projects such as the Human Microbiome Project(*9*) and MetaHIT(*10*, *11*), which have focused on particular niches within the body. However, our closest genetic neighbours, non-human primates (NHPs), have not received nearly as much attention by comparably large scale microbiome studies which tend to focus on only a few NHP species(*12*), target specific pathogens(*13*, *14*) or use 16S rRNA(*15*) approaches which don’t have the ability to discover new viruses.

In this study, we assembled and annotated non-host sequences derived from cell-free DNA (cfDNA) from blood samples of of 221 individuals (322 samples) from an assortment of both Great Apes (two species) and Old World Monkeys (15 species) (Fig. 1A) from three wildlife refuges in the West African nation of Cameroon (Fig. 1B). The breadth of taxa present in these animals provides a resource of the baseline microbiome for numerous previously unstudied species. Combining our molecular data with the geographic sampling shows that the environment plays a similar role as the evolutionary history of a given species role in shaping the total microbiome, but that (non-phage) viruses have a stronger host and species association. We found evidence of multiple potential zoonoses, including a potential four-level infectious relationship: primate - amoeba - virus - virophage, with the virus being a large nucleocytoplasmic large DNA virus most similar to African Swine Fever Virus, new strains of the Hepatitis B virus that appear to have the ability to infect multiple species, and reservoirs of pathogens including *Treponema pallidum*, the causative agent of diseases such as syphilis, yaws and bejel. Taken together, our results provide a wealth of information about both bacterial and viral diversity in NHPs.

**Figure 1:**
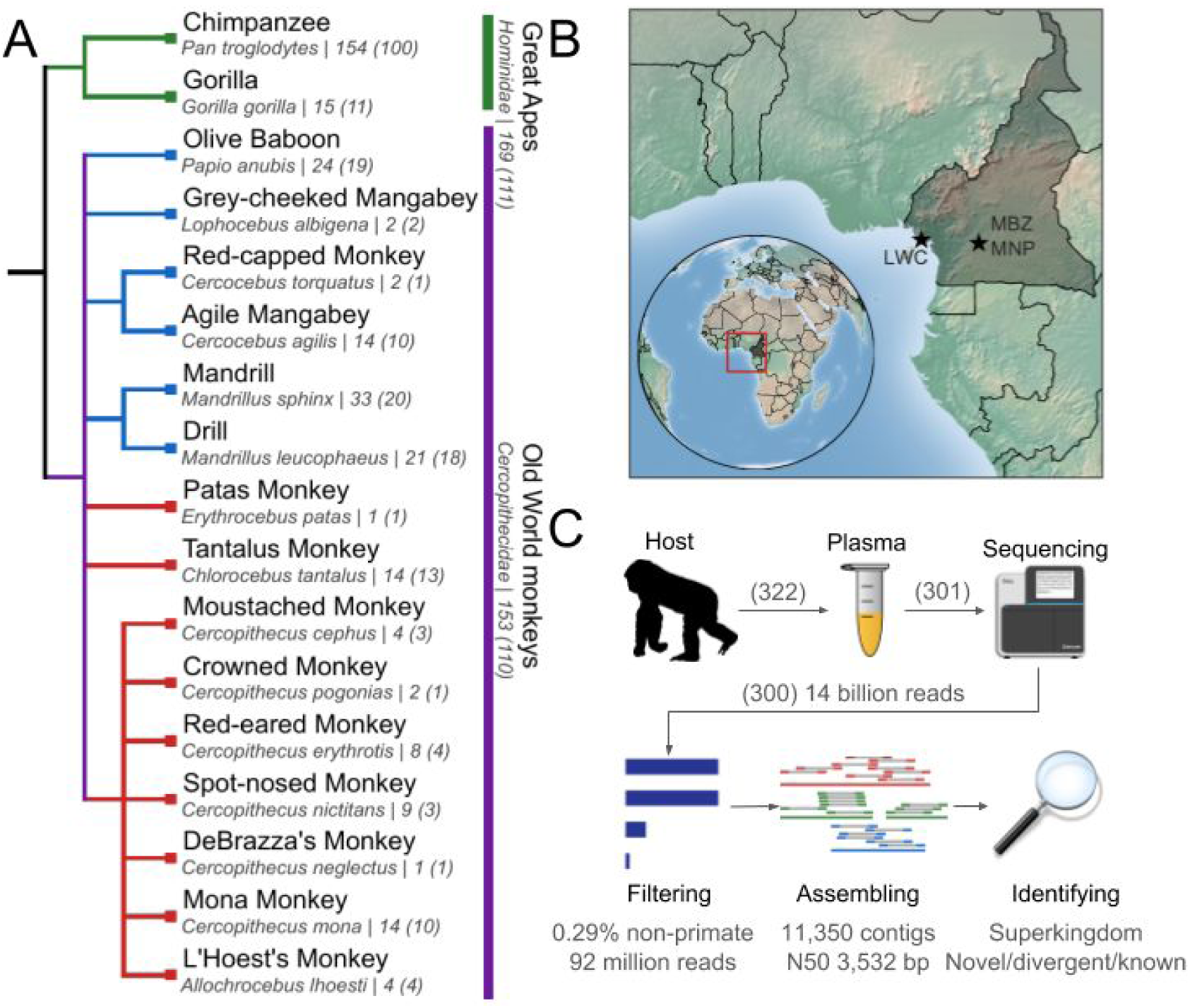
Overview of sample origin and processing. A) Phylogenetic tree of the species sampled in this study. Levels shown (from left to right) are order (primate), family (Hominidae - green, Cercopithecidae - purple), genus and species (common and scientific name, number of samples (individuals) indicated by numbers after the vertical line). In Cercopithecidae, the tribes Papionini (top, blue) and Cercopithecini (bottom, red) and are shown. B) Map of Cameroon, a West African country showing the locations of the three locations from which samples were sourced. Key: LWC - Limbe Wildlife Centre, MBZ - Mvog Betsi Zoo, MNP - Mefou National Park C) Simplified illustration of the sample processing pipeline. Plasma obtained from a primate has circulating nucleic acids extracted, sequenced, filtered for non-primate reads, assembled and the potential superkingdom and novelty is determined. Numbers in parentheses are the number of samples used after each step. Quantitative details in Fig. S2 and S3.

Samples were obtained over three years, with 70% of individuals providing only one sample (Fig. S1). The animals resided in three wildlife sites in Cameroon: the Limbe Wildlife Centre, Mvog Betsi Zoo and Mefou National Park (the last two under the management of Ape Action Africa, formally the Cameroon Wildlife Aid Fund). After plasma was obtained, 301 samples had sufficient quality or quantity of DNA to undergo high throughput sequencing (2×75bp, Illumina NextSeq 500), resulting in a total of 14 billion reads (2.1 trillion bp), with a median of 45 million reads per sample. Three samples were found to be mostly contaminants (low proportion of primate sequences) and excluded from further analysis, and two samples were sequenced twice, resulting in a total of 300 samples (Fig. 1C, upper). Multiple quality control measures were applied to the reads to remove low quality bases, merge overlapping reads (from short fragments) and to remove various biological vector sequences (Fig. 1C, lower). These processes removed only a small proportion of reads (Fig. S2B-D). Finally, the reads were aligned to a suitable reference primate genome, after which the nonhost reads (median of 1.2%) were collected and then aligned sequentially to databases of a further twelve reference primate genomes as well as the NCBI nucleotide database (Fig. S2G-H), resulting in the removal of an additional 70% of reads. This resulted in a median of 0.29% of reads being classified as nonhost, for a total of 92 million reads used in subsequent analysis. This proportion of nonhost reads is comparable to our observations in similarly processed human samples.

Non-primate reads were assembled in two stages: first by individual and second by species. In both stages, genes were predicted and nucleotide sequences were tested for homology. Reads aligning to the assembled contigs found to be of a primate origin or marked as low complexity were removed. Across all species assemblies, a total of 33,853 contigs longer than 1 kbp were produced. A series of filtering steps was undertaken to deplete potential contaminants and enrich for informative sequences (Fig. S3A). First, only contigs with at least 60% gene coverage were kept (removing 2,670 contigs). Then, using a set of six sequencing libraries (a total of 275 million reads) prepared using the same protocol for cfDNA extraction from plasma, but using water or DNA from a human cell line, we removed an additional 5,745 contigs that had at least 10 reads/kilobase of sequence aligning to them. Finally, to conservatively retain only contigs that are unlikely to be kit contaminants, an additional 14,088 contigs were removed which shared a taxonomic assignment with the potential contig contaminants with high levels of homology. Over two thirds of the ‘contaminant’ contigs were Pseudomonas variants, with many examples of Shewanella, Bacillus, Achromobacter and Saccharopolyspora as well. This resulted in a final set of 11,350 contigs, with an N50 of 3,532 bp (Fig. S3B).

Contigs were taxonomically classified based on the consensus taxonomy of gene homologies. They were further partitioned into known, divergent and novel contigs (Fig. S3). Known (red) means that the BLAST coverage was at least 20% and average gene identity above 80%. Novel (green) means that both BLAST coverage and average gene identity was below 20%, and divergent (yellow) is everything in between. The breakdown of taxonomic level to which contigs are assigned to is presented in Fig. S3D. Many of the shortest contigs (those with only a single gene) were novel contigs, which while having low gene homology, were by default assigned to the species level. Otherwise, the overall trend is for novel contigs to be assigned to higher taxonomic levels. This can be seen qualitatively in the solar system plots (Fig. S4), where the novel contigs clearly have a high mass of contigs assigned to the root of the taxonomic tree (yellow dots in the outer orbit). Most (n=8143) of the contigs appear to be bacterial in nature, with a large number assigned to the root (n=1999), hundreds assigned as eukaryotes or viruses and a few (n=46) as archaea (Fig. S3E and S5). Tables of both the counts of reads aligning to contigs, those contigs annotations and tables of the taxid present in all samples are available in the supplementary datasets.

A comparison with our previous novel human microbiome data(*8*) was performed by comparing the assembled NHP-derived contigs to those ones using LAST. Of the 11,350 NHP contigs, 884 had good matches (50% alignment of contigs at least 1 kbp in length). The majority of these were novel (n=472) with the remainder (n=412) as divergent. Most were bacteria (n=619), such as various Proteobacteria and the matched viruses (n=73) were all phage-like. These contigs lay in 347 taxids, which are a subset of the 2120 taxids of the remaining NHP contigs that did not have homology to the novel human contigs. Comparing the presence of these contigs in the human and NHP samples (Fig. S6), at the low level there is a clear clustering by study, but with an overall structure of mixing the human and non-human samples together, indicating a high level of sharing of taxonomic features across the primates.

Many of these taxa have been previously identified in the microbiome of humans and NHPs (annotated taxa and relative abundances shown in Fig. S7). Of the bacteria, many contigs were assigned to families such as Flavobacteriaceae, Pseudomonadaceae, Yersiniaceae, Enterobacteriaceae, Caulobacteraceae, Burkholderiaceae and Bacillaceae. Archaea included representatives from the phyla Crenarchaeota, Euryarchaeota and Thaumarchaeota, including the orders Methanobacteriales, Sulfobales, Thermoplasmatales, Halobactericale and Nitrosphaerales that have been observed in human microbiomes before(*16*, *17*). Besides a small number of eukaryotic contigs that appear to be chordata (and hence likely to be from the host), multiple contigs were assigned to the Apicomplexa phylum, which includes the parasites: *Plasmodium* that causes malaria, and *Babesia* that causes babesiosis. In addition, many fungi including known pathogens in the Tremellales order (such as *cryptococcus*) and fungal orders containing highly prevalent genera(*17*, *18*) such as *Candida*, *Cladosporium*, *Aureobasidium* and *Saccharomycetales* also had novel and divergent contigs. In addition, amongst the eukaryotic contigs were numerous plants, likely derived from the diet(*19*).

The majority of viral families (7/12) known to infect primates were found in the *de novo* assemblies including: adenoviridae, anelloviridae, hepadnaviridae, herpesviridae, parvoviridae, polyomaviridae and retroviridae. However, of the assembled viruses, most were phages (Caudovirales or Microviridae).

Multiple individuals had long (>1 kbp) hepatitis B virus derived contigs, so in order to better describe their prevalence and evolution, 11 reference guided consensus sequences (labeled A-K) were found and a phylogenetic tree was produced using sequences from human and other non-human primate strains. The overall phylogenetic structure (Fig. S8A) recapitulated the known NHP HBV genotypes(*20* – *23*), in that there are three chief branches: human sequences (genotypes A-G clustering within themselves, and the two NHP samples interleaved were known from the literature(*23*, *24*)); Asian NHP sequences (chimpanzee sequence also known from the literature); and African NHP sequences, composed of gorilla and chimpanzee sequences, along with our sequences deriving from various species. Our sequences lie within the existing chimpanzee and gorilla genotypes(*25*) with high confidence (Fig. S8B).

Thirty-nine individuals across ten species were observed with HBV (Fig. 2A and S8C) in Mefou National Park and the Limbe Wildlife Centre. Across both parks, chimpanzees were identified with chimpanzee HBV genotypes. In contrast, the gorilla HBV lineage was present in both gorillas and chimpanzees in the Limbe Wildlife Centre. Many Old World monkeys in this park were also found to have gorilla lineages present, although some have low coverage that can’t disambiguate between the two lineages. In particular, this study provides the first sequence based evidence that eight species in the Papionini and Cercopithecini tribes can be infected with HBV. The ability to observe high coverage of viral diseases such as HBV while providing new lineages and infection patterns demonstrates a benefit of applying an untargeted approach to microbiome sequencing.

**Figure 2:**
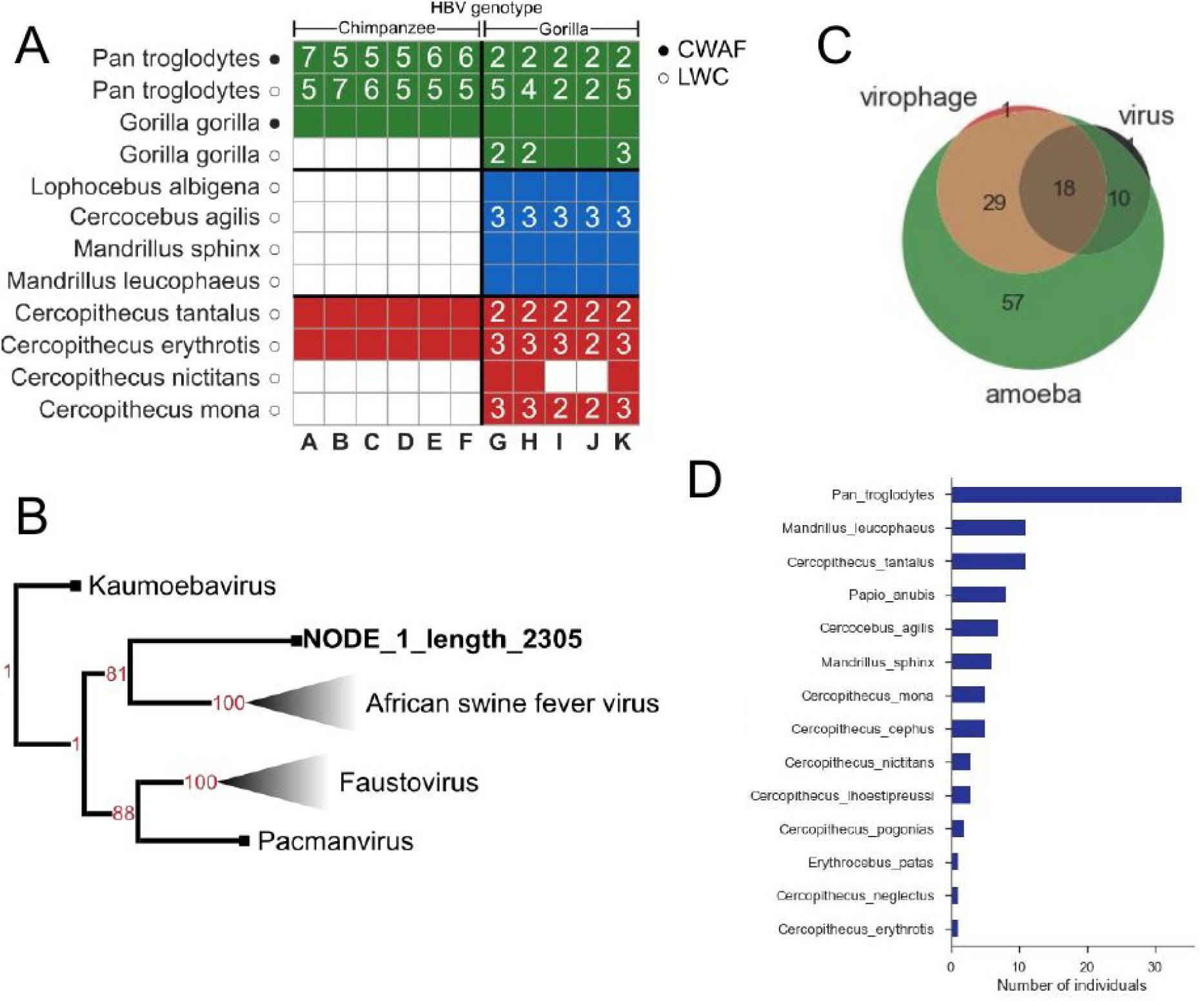
Details of three potential zoonotic diseases. A) Heatmap showing the HBV genotypes (A-F, chimpanzee lineage; G-K, gorilla lineage) with support in different species stratified by park. Filled squares without numbers have a value of “1”. B) Tree (with bootstrap in red) of other NCLDVs with homology to the gene found on the longer of the contigs. Faded triangles indicate multiple sequences below that point. C) Venn diagram of the number of individuals found to have amoeba, NCLDV and associated virophage sequences. D) Number of individuals in each species with evidence for Treponema pallidum bacteria.

Two contigs had low homology hits to a polymerase from African Swine Fever Virus (ASFV), an arthropod borne dsDNA virus that causes haemorrhagic fevers in domesticated swine(*26*). By recruiting additional reads and and reassembling, two contigs were produced, the larger being 2,305 bp, the shorter 1,386 bp. Genes were predicted, and a database of the nucleocytoplasmic large DNA viruses (NCLDVs) was used to find homologous sequences, generate a multiple sequence alignment build a phylogenetic tree (Fig. 2B and supplementary Fig. 9A-B). These contigs are a likely sister group of the clade including ASFV and the Faustoviruses(*27*) and Pacmanvirus(*28*). The high bootstrap values and concordance of the rest of the (reference) NCLDV tree with the literature leads confidence in this placement. ASFV is known to infect pigs, but many of the other NCLDV infect amoebas(*29*). This led to identifying candidate amoeba contigs (n=10), as well as virophages (n=2). These were found to mostly coinfect the hosts (Fig. 2C and S9C), suggesting that cfDNA analysis has picked up a four level infection: primate, amoeba, eukaryote-infecting virus and virophage.

The high-throughput hypothesis-free sequencing approach also allows for monitoring of certain diseases known to have reservoirs in NHPs. *Treponema pallidum(30*, *31)*, is one such species that includes subspecies that cause the human diseases: syphilis (ssp pallidum), bejel (ssp endomicum), pinta (ssp carateum) and yaws (ssp pertenue). By creating a database of known strains and aligning the reads to these, 98 of the non-human primates were found to have reads aligning to this bacteria, although coverage was too low to disambiguate effectively between the different strains (Fig. 2D). Nonetheless, it provides further evidence of the need to monitor non-human primates as human disease reservoirs(*32*).

We searched for sequences that could be part of the candidate phyla radiation (CPR)(*33*) in the non-human primate data. This encompassed downloading sequences from the NCBI nucleotide collection (they were not in the BLAST database) and aligning all contigs to this sequence. Those that matched potential CPR genomes were extracted and then aligned to the nt database. Finally, the scores of these alignments were compared, resulting in 62 contigs being found that best align to CPR contigs. Therefore, like in humans, CPR is represented in the NHP microbiome, perhaps originating in the drinking water.

The large number of species in this dataset and the fact that they originate from three locations allows us to investigate the extent to which the structure of the microbiome is due to the evolutionary history of the organism (i.e. cospeciation), compared with the influence of the environment or diet(*34*, *35*). To account for the absolute differences in the number of nonhost reads, counts were rarefied to allow for comparisons between samples. Comparisons between samples was made in a taxonomically informed manner, using the average of the unweighted UniFrac distance calculated for different ranks. We used different ranks to allow the less-confident assignment of novel and many divergent contigs to still play a role in determining taxonomic similarity between samples, which would have been missed if only a single (e.g. species or genus) rank was used.

Distributions of distances were computed for all samples, within individuals and in all combinations of restricting to the same or different park or species (Fig. S10A and S11A). Differences of empirical distribution functions was computed pairwise for all such comparisons (equivalent to the Kolmogorov-Smirnov statistic) and the region found to be significantly different (p < 0.01) was found. When all taxa were included, the hierarchy of influences on the microbiome was found to be dominated by species in the same park (purple line). These samples are much more similar to each other than those from just the same park or just the same species (red and blue lines). Although the influence of either park or species is not found to be strong, when the other condition is different (dotted lines), the UniFrac distance between samples increases (i.e. they are more dissimilar taxonomically). When restricted to non-phage viruses, individuals (Fig. S11A) and species (Fig. 3B) have a much stronger role in determining whether viromes are similar.

**Figure 3:**
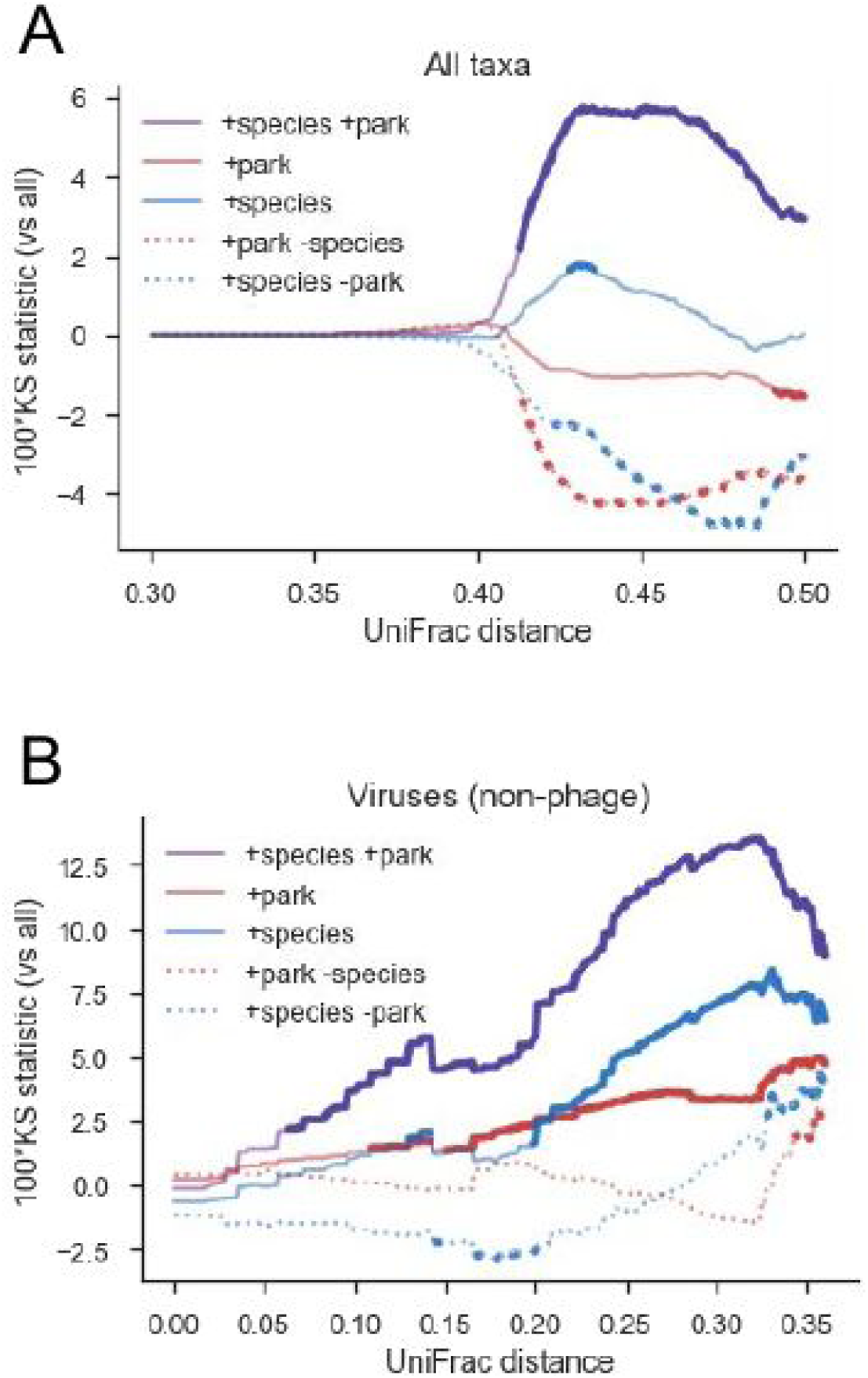
Difference in empirical distribution function of multilevel UniFrac distances between certain classes of samples. A) All taxa used in computing the UniFrac distances. Plots show 100x the Kolmogorov-Smirnov statistic for subsets of samples restricted to being from the same (+, solid)or different (-, dotted) park or species. Thicker lines indicate a significant difference in distribution compared to all (p < 0.01). B) Same as A), but taxa were restricted to non-phage viruses. Supplementary figures 8-9 show details of all comparisons and a graphical representation of sample distances.

This result can also be visualized by inspecting the tSNE projection of the UniFrac distances (Fig. S10B and S11B). For a given host species, the samples tend to be not cluster tightly when all taxa were used, with parks partitioning the samples completely for *Cercocebus agilis* and *Cercopithecus tantalus*. In contrast, when viruses are investigated, species tend to cluster in smaller areas associated with their tribe, and the parks have much more overlap. The conclusion of this is that the cellular microbiome is highly transferable amongst primate species co-located in the same park, but viruses show higher levels of host specificity.

Deep sequencing of circulating nucleic acids continues to reveal the remarkable result that the majority of non-host sequences in primates are novel or divergent, with the power to recapitulate the known component in an unbiased manner. The ability to identify the majority of primate infecting viruses without targeted assays demonstrates the potential for discovering new viral pathogens using this method. Given the high level of sharing of the microbiome across primates, as evidenced by the strong environmental influence on its structure, with further deep sequencing of primates and expanded unbiased assays of circulating nucleic acids, we expect to be able to identify potential zoonoses before a spillover is detected, as well as better understand the basic structure of our microbiome.

## Acknowledgments

We would like to thank the Government of Cameroon, Limbe Wildlife Centre, Mefou National Park and Mvog Betsi Zoo for authorisation to collect and export samples. We would also like to thank the staff of Limbe Wildlife Centre, Mefou National Park and Mvog Betsi Zoo who assisted in collection of samples and the staff of the Military Health Research Centre, Yaounde (CRESAR) and Metabiota Cameroon who assisted in the management of the collections and shipping. Funding: This work benefited from support from the United States Agency for International Development (USAID) Emerging Pandemic Threats PREDICT program (Cooperative Agreement no. AID-OAA-A-14-00102). This work was supported by the John Templeton Foundation as part of the Boundaries of Life Initiative (Grant 51250 and 60973). Author contributions: MK, ML, NDW and SRQ wrote the manuscript; IDV, JO and NFN processed samples; ML, JN, UT, JDL, BT and JK devised sampling protocols, collected samples and managed collections and shipping. Competing interests: Authors declare no competing interests. Data and materials availability: Raw sequencing files are available on the NCBI SRA under accession numbers …. Assemblies, annotations and trees are available in the supplementary material. Bioinformatics pipeline is available at code.stanford.edu …

## Supplementary Materials

### Materials and Methods

Plasma was extracted from whole-blood samples from the non-human primates in the same manner as previously described(*1*) and stored at -80C. Samples were later thawed and cell-free DNA was extracted using the QIAmp Circulating Nucleic Acid Kit (Qiagen). Libraries were prepared using an automated microfluidics based platform (Mondrian ST; Ovation SP Ultralow Library Systems). Libraries were quantified on an Agilent 2100 Bioanalyzer (High Sensitivity DNA Kit) and sequenced on an Illumina NextSeq 500, in 2×75bp high output mode (∼45 million reads/sample, 8 samples/library).

Demultiplexed reads were then processed in a custom bioinformatics Snakemake(*36*) pipeline. Firstly, read quality was assessed by FastQC(*37*); low quality bases and adapter sequences were trimmed using Trimmomatic(*38*); mate pairs that overlap were merged using FLASH(*39*); various linkers, primers, other adapter and vector sequences commonly used in labs were removed by aligning reads using bowtie2(*40*) against the UniVec(*41*) core database. This resulted in a set of ‘clean’ reads.

These reads were then aligned using bowtie2 to each of twelve different primate reference genomes as downloaded from ENSEMBL. These are: bushbaby, chimpanzee, gibbon, gorilla, human, macaque, marmoset, mouse lemur, olive baboon, orangutan, tarsier and vervet. Reads that didn’t align were extracted. An additional set of alignments was performed sequentially after aligning to the “best” reference for each species (based on the tribe, see figure 1A) in this study (supplementary table below). Following alignment to all 12 primates, to capture additional primate sequences we aligned using BLAST(*42*) (blastn, megablast mode, percentage identity >80%, evalue < 1e-4) against NCBI’s nucleotide database and filtered Eukaryotic matching read with any of the following in their name “human, chimpanzee, gorilla, orangutan, gibbon, macaque, lemur, monkey, Cercopithecus, grivet, Barbary, baboon, marmoset, mangabey, tarsier, lutung, tamarin, guereza, siamang”. This resulted in a set of non-host reads.

| Species | Reference genome |
| --- | --- |
| Pan troglodytes | chimpanzee |
| Cercopithecus erythrotis | vervet |
| Gorilla gorilla | gorilla |
| Lophocebus albigena | olive baboon |
| Cercopithecus mona | vervet |
| Mandrillus sphinx | olive baboon |
| Mandrillus leucophaeus | olive baboon |
| Cercopithecus tantalus | vervet |
| Cercopithecus nictitans | vervet |
| Cercopithecus cephus | vervet |
| Cercopithecus pogonias | vervet |
| Cercopithecus neglectus | vervet |
| Erythrocebus patas | vervet |
| Cercocebus agilis | olive baboon |
| Cercocebus torquatus | olive baboon |
| Papio anubis | olive baboon |

The non-host reads for each individual were used as the inputs of the first set of assemblies. They were assembled using SPADes (version 3.5.0)(*43*) using the default settings. Following assembly, contigs were BLASTed (task=blastn) to the nt database and low complexity sequences found using dustmasker(*44*). Contigs were marked as ‘bad’ if they were low complexity over 75% of their bases or the primate-like reads (using the same name filters as above, but removing those contigs that also aligned to non-Eukaryotic sequences). Non-host reads from each individual were aligned to the assembled contigs, and reads that aligned to the bad contigs were filtered out.

These reads were then used as the input for the next set of assemblies, grouped by the species of origin. A similar assembly process and post-assembly alignment and cleanup was performed. In addition, genes were predicted using PRODIGAL(*45*) and predicted gene sequences aligned for homology (blastx, NCBI nr database). To determine homology between assembled contigs, all assemblies (individuals and species) were aligned to each other using LAST(*46*).

To determine viral families known to infect primates, of the 7,431 complete viruses on NCBI, the 1,613 with vertebrate hosts were selected. These corresponded to 1,429 entries in Genbank which were downloaded, and after querying the host entry, 287 had primate hosts. Selecting for DNA viruses resulted in 146, which were then classified based on their ICTV family. The hepadnaviruses were also added to the list of primate infecting viruses.

Microbiome reads for each sample were rarefied (subsampling to 1000 reads, n=100 repeats) and all taxonomic levels identified recorded to a file. Then, a taxonomic tree built using the NCBI reference taxonomy for all identified taxa was built. For each taxonomic level (superkingdom, phylum, class, order, family, genus and species), the total tree is pruned to keep only leaves of at least the given level. The unweighted UniFrac distance is then computed pairwise for each sample. The average of the UniFrac distances is taken over the different taxonomic levels, and then the median over the rarefied samples taken. For the taxonomically restricted comparisons, the taxa present in a sample were required to be a virus (taxid: 10239) but not a phage (taxids: 28883, 79205, 38018, 10841, 12333, 552364).

Distributions of distances were computed for all samples, within individuals and in all combinations of restricting to the same or different park or species. An empirical distribution function was found, and differences between sets of the populations computed (Kolmogorov-Smirnov statistic). Visualization of the samples was performed using tSNE on the median level-average UniFrac distances. Ellipses indicate the region of select samples and were determined using a minimal volume enclosing ellipse algorithm(*47*)

Contigs that were assigned to the taxid 137992 (Asfarviridae) and those that had local homology to these were chosen from the assemblies, resulting in 10 contigs. A bowtie2 database was made from these, and for each sample, the non-primate reads were aligned to this database. The aligned reads (171 paired end and 70 unpaired reads) were extracted, and SPADes used to assemble the reads. This resulted in three contigs, the two largest of which kept the Asfarviridae homology and were retained. A similar process of building a database based on these contigs, recruiting reads and assembly was repeated 8 more times, until a steady state resulted (the largest contig increased in size by over a 1 kbp, 180 paired end and 85 unpaired reads recruited). Genes were predicted with PRODIGAL, and a database of the nucleocytoplasmic large DNA viruses (NCLDVs) from NCBI (taxids: Ascoviridae - 43682; Asfaviridae - 137992; Iridoviridae - 10486; Marseilleviridae - 944644; Mimiviridae - 549779; Phycodnaviridae - 10501; Pithoviridae- 2023203; Poxviridae - 10240; Dinodnavirus - 1232647; Faustovirus - 1477405; Kaumoebavirus - 1859492; Pacmanvirus - 1932881) built and regions homologous to the predicted gene found on these contigs and extracted.

All these sequences were aligned with MUSCLE(*48*) using the default settings. A phylogenetic tree was produced with RAxML (version 8.2.10)(*49*), using the GTR+CAT model and bootstrapping 100 samples. The tree was imported into python using ete3(*50*) for visualization and annotations.

Counts were determined from the reads recruited. The additional amoeba contigs were found by selecting for taxid 554915. The virophage ones for taxid 552364. Counts for these contigs were determined from the previously created count matrix.

Non-host reads from all samples were aligned against reference primate HBV sequences (accessions: JQ664502, JQ664505, JQ664506) from an existing study of some of the same individuals(*25*) using bowtie2 (--local mode). Aligned reads were sorted and multiple pileups (samtools mpileup) created for each individual. Consensus sequences were found using varscan (version 2.4.2, ‘mpileup2cns’ command)(*51*) and VCFtools(*52*). For each individual, the chosen consensus sequence was determined by which of the three references had the highest coverage. These sequences were compared and merged into a set of 11 unique sequences.

To augment these sequences with other non-human primate HBV sequences, the NCBI nucleotide database was searched for genomes between 3000-3300 bp and having the taxid 10407. This resulted in 9466 Genbank entries being downloaded. These were filtered for entries that had non-human hosts, resulting in 89 sequences. Human sequences were downloaded from HBVdb(*53*) with three exemplar sequences from each of eight human genotypes (A-G) being selected. Additionally, two sequences from woolly monkeys(*54*) were included as an outgroup. In total 126 sequences HBV sequences were used, reducing down to 123 once accounting for duplicates.

Like for the NCLDVs, these sequences were aligned with MUSCLE(*48*) using the default settings and a phylogenetic tree was produced with RAxML (version 8.2.10)(*49*), using the GTR+CAT model and bootstrapping 100 samples, this time specifying the woolly monkey sequences as the outgroup.

Reference genomes were taken from NCBI by searching for sequences assigned to the taxid 157. All non-primate reads were aligned to each of these. These were grouped together by species and those reads that aligned to *T*. *pallidum* and not to other species in the genus were kept. These reads were then summarized by individual, resulting in a table of *T*. *pallidum* detection that can be aggregated by species. Attempts were made to disambiguate between the subspecies, however coverage nor SNV presence was not high enough.

Figures 1-3

## Figures S1-S11

**Supplementary figure 1:**
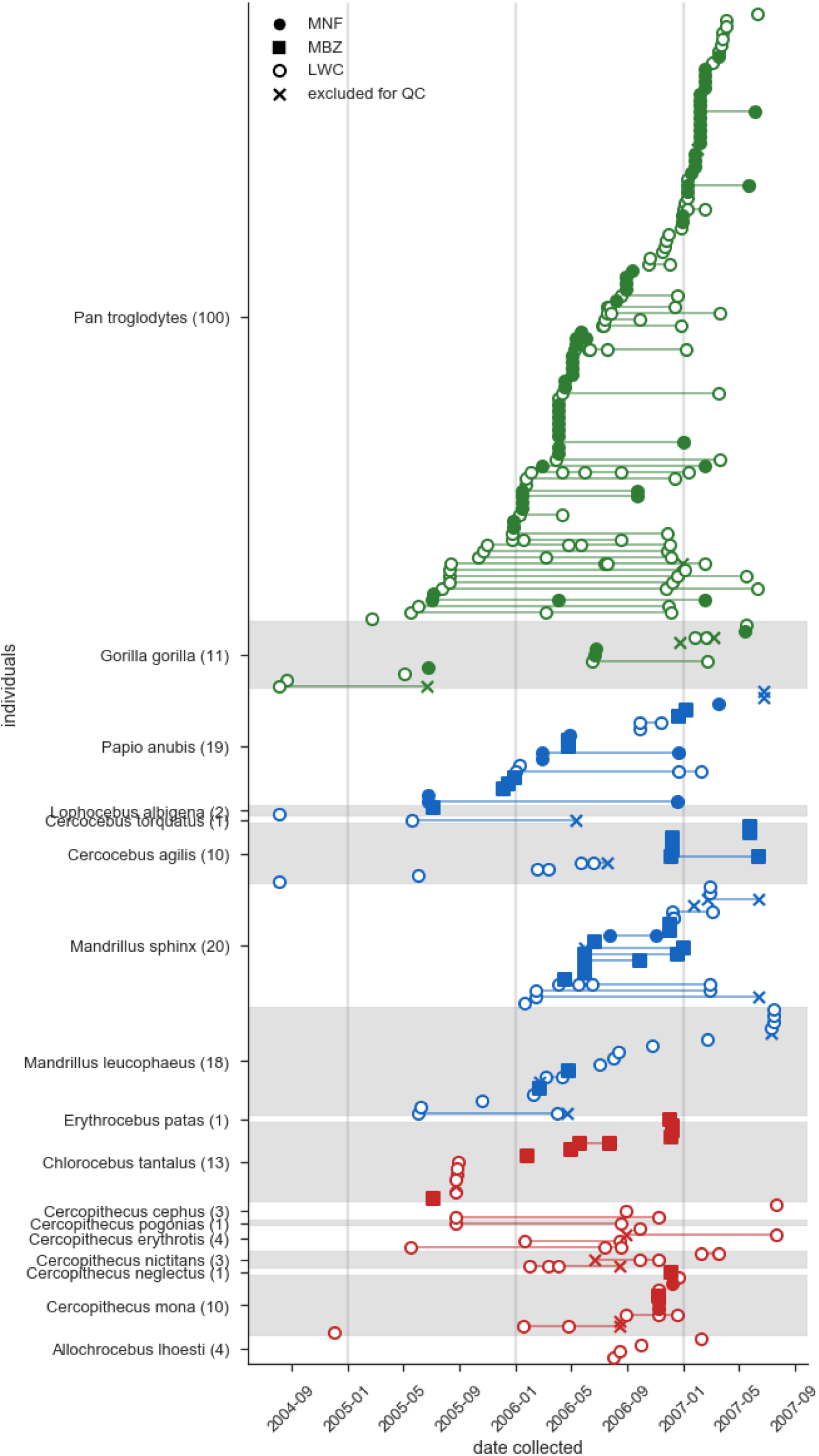
Timeline of sample collection. Species are stratified in the same order as the phylogenetic tree of figure 1A. Samples within each species are ordered by first time sampled for each individual. Points are labelled by the park they were from and whether they were sequenced successfully (circle/square - success; cross - failure).

**Supplementary figure 2:**
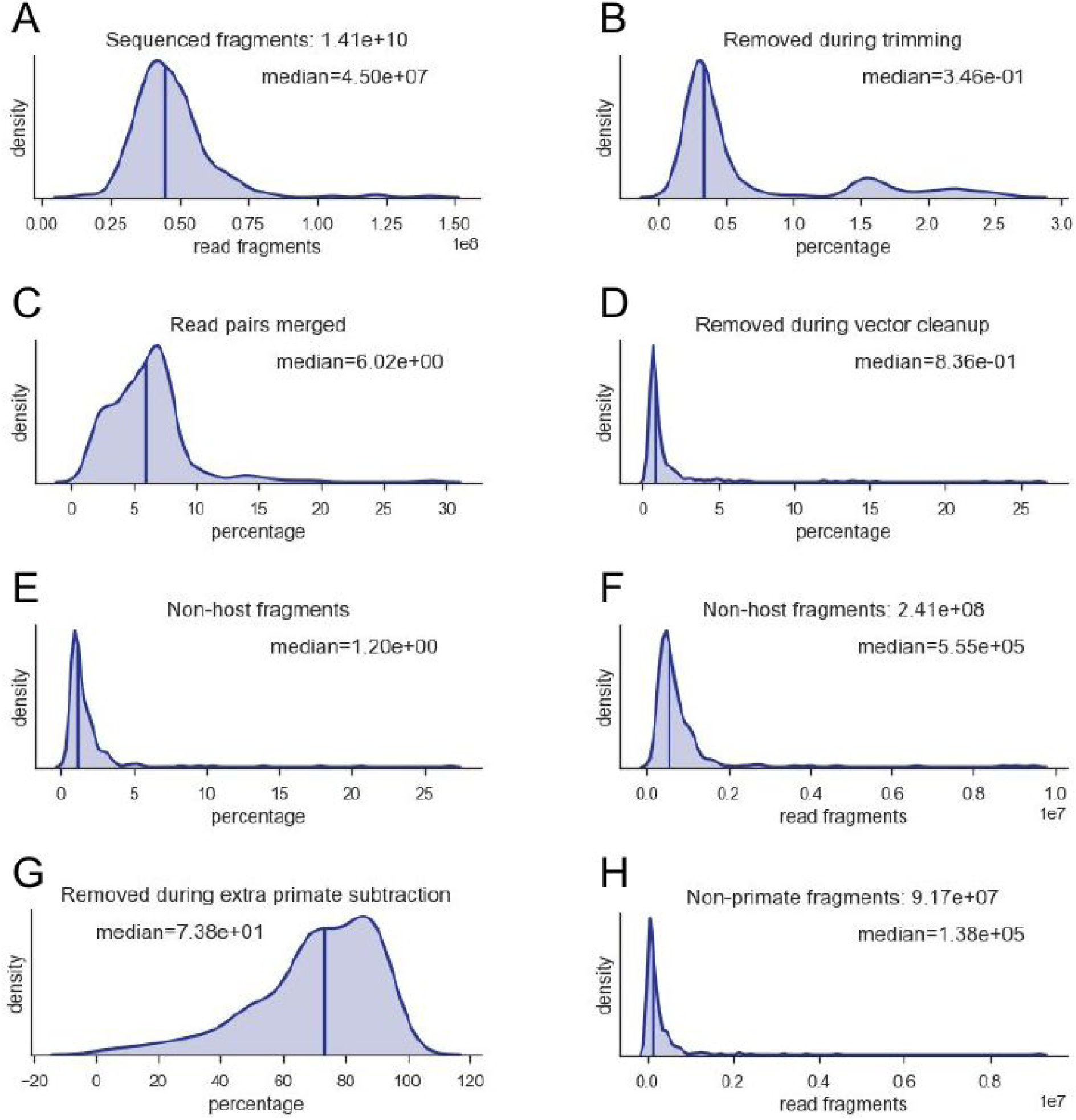
Sequencing and processing pipeline distributions. Vertical lines indicate the median value annotated on plot. A) Raw read fragments (2×75 bp) sequenced. B) Percentage removed during trimming. Percentage of reads merged. D) Percentage removed during vector cleanup. E) Percentage of cleaned reads that do not align to the closest host genome. F) Number of reads that remain after host removal. G) Proportion of non-reference host reads removed by additional alignment steps to other primate genomes. H) Number of reads remaining that are of a non-primate origin.

**Supplementary figure 3:**
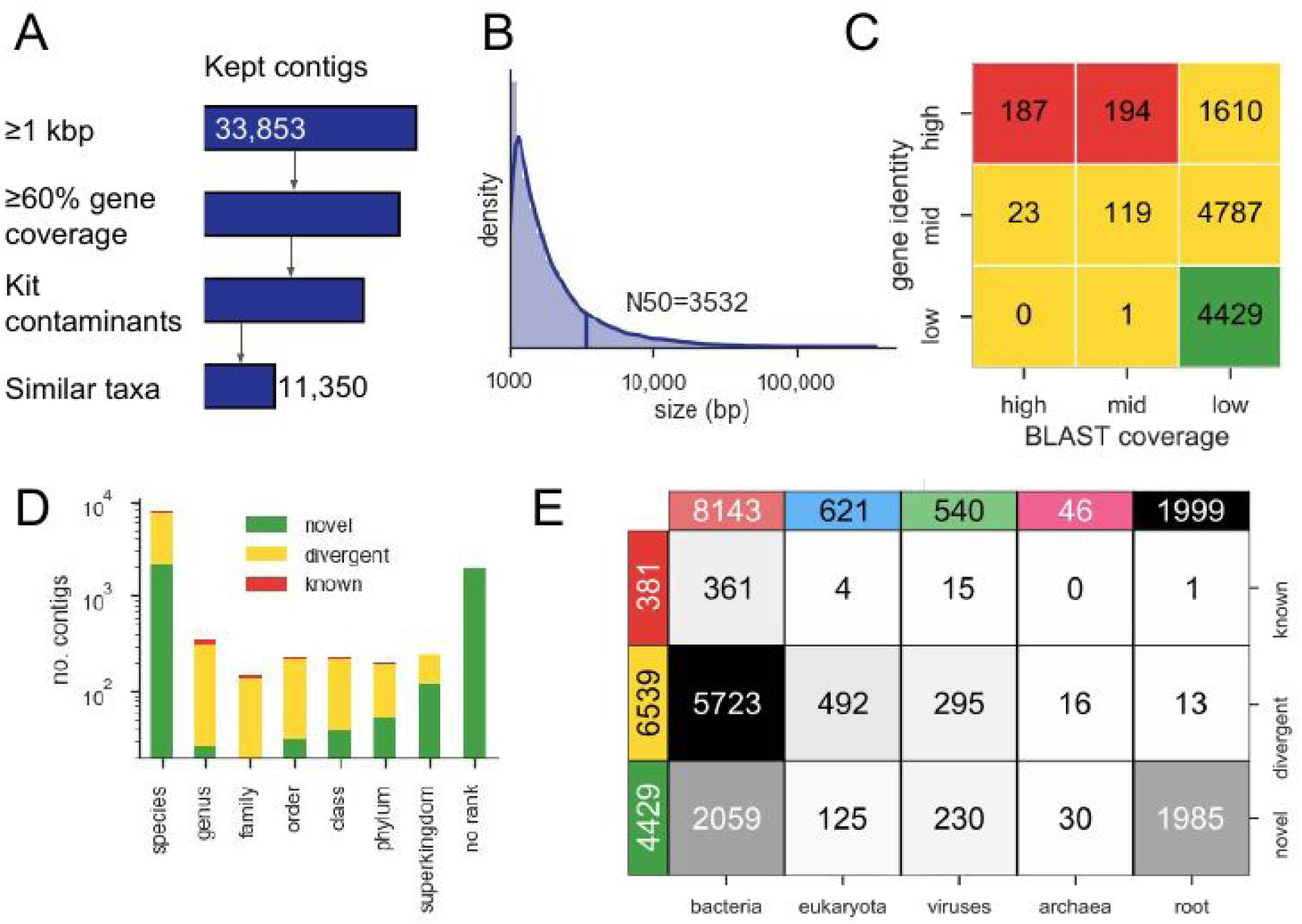
Description of the assembled contigs. A) Filtering pipeline of the contigs. 33,853 contigs were at least 1 kbp. Those with above 60% gene coverage were kept. Contigs that had at more than 10 reads/kilobase aligning to them across all the control samples were removed as potential kit contaminants. Finally, contigs sharing any of the contaminant taxa on known contigs were removed, leaving a final set of 11,350. B) Size distribution of the selected contigs. C) Number of contigs falling into the high/mid/low gene identity and BLAST coverage. Colours signify novel (green), divergent (yellow) and known (red) contigs. D) Stacked barchart of the number of contigs assigned to different taxonomic ranks. E) Annotated heatmap of the number of contigs assigned to each superkingdom stratified by novel, divergent and known contigs.

**Supplementary figure 4:**
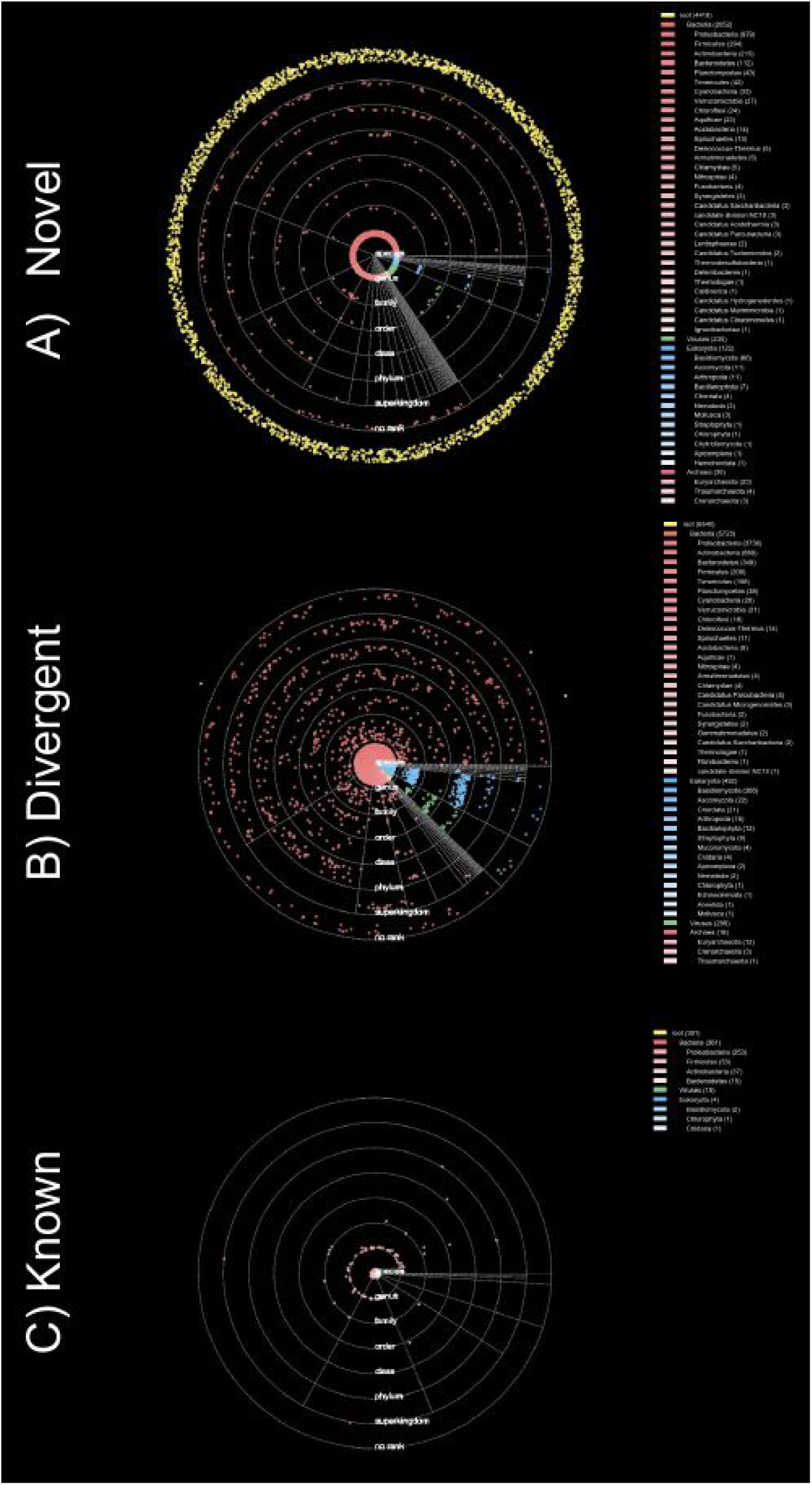
Solar system plots. Each dot corresponds to a contig, colour based on the superkingdom, saturation based on the phylum. Each ring represents a taxonomic level (see figure S3D), with the intraring radius proportional to the average gene homology. The unrooted contigs (yellow dots) outside the last ring could not be assigned to any superkingdom; the radial scattering is to help illustrate the density of contigs. Legend on the right of each solar system plot is a tree showing the number of contigs assigned to each superkingdom and phylum present. A) novel B) divergent C) known.

**Supplementary figure 5:**
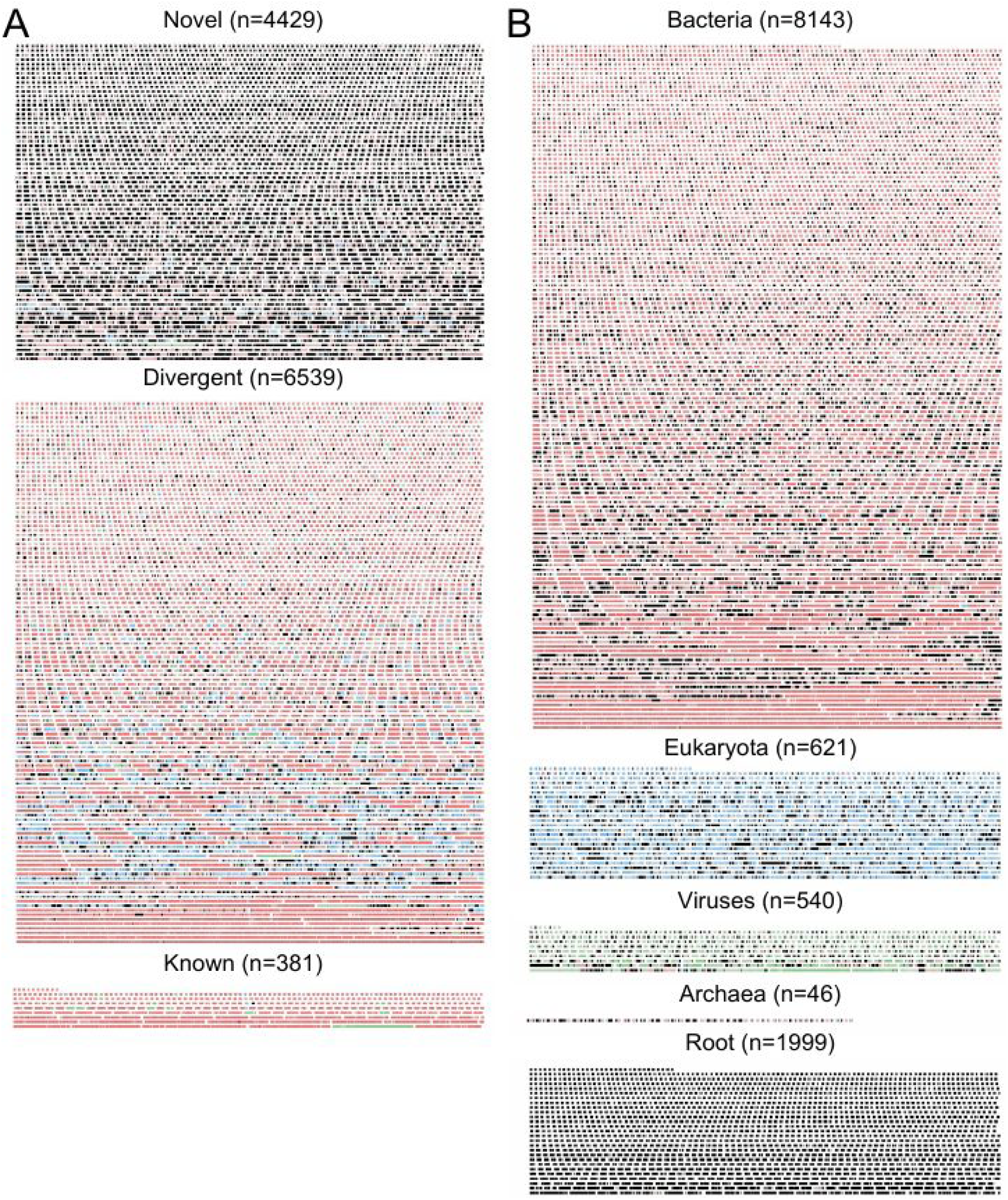
Contig packing plots. Each rectangle is an illustration of a contig, the width of the whole block is 200 kbp. Genes on each contig are colored by the superkingdom they are most similar to, saturation proportional to gene identity. A) Contigs partitioned as novel, divergent and known. B) Contigs partitioned to superkingdom.

**Supplementary figure 6:**
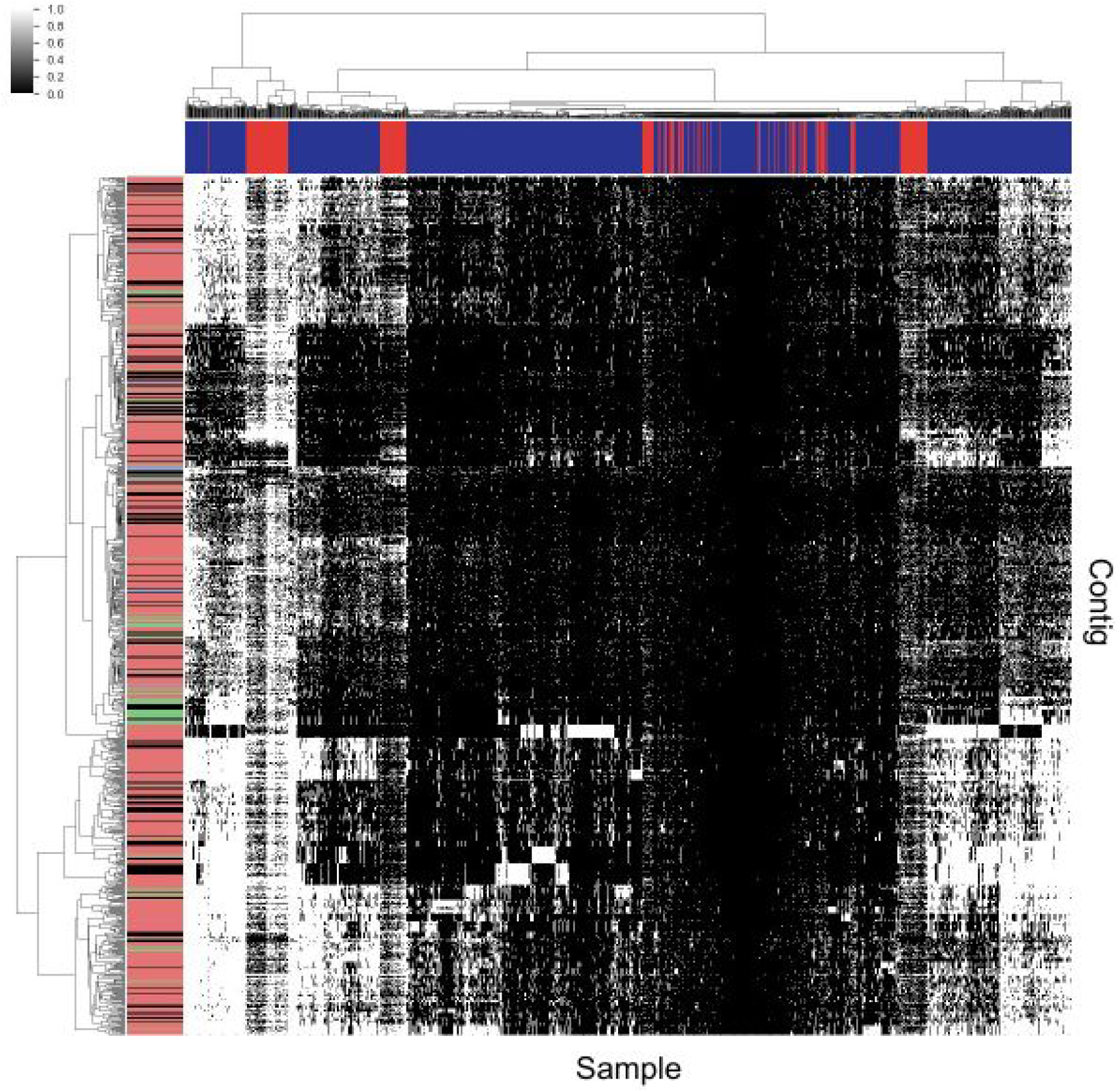
Contig presence in human and non-human primate samples. Colors for the columns are red (NHP) and blue (human). Dendrograms are optimally ordered and clustered using Ward’s method on a Hamming distance matrix. Rows are the 884 contigs found to be shared between these assemblies and previous ones derived from humans.

**Supplementary figure 7:**
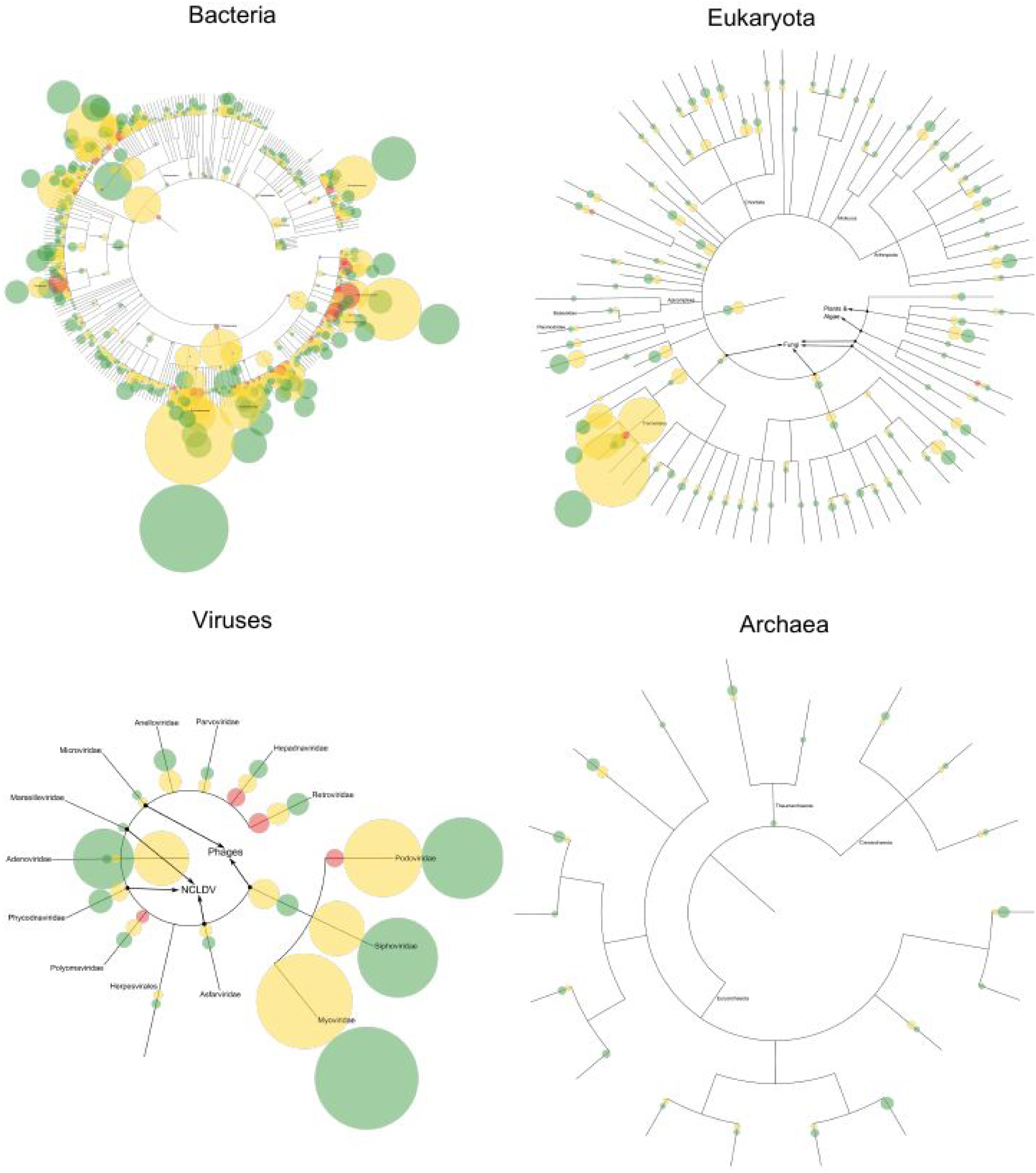
Selectively annotated taxonomic trees based on the assembled contigs. Each tree is based on the NCBI taxonomy for contigs assigned to each of the four superkingdoms. Circles are proportional to the number of contigs, each colored by whether they are known, novel or divergent. Full annotated trees are available in the supplementary material in the Newick format.

**Supplementary figure 8:**
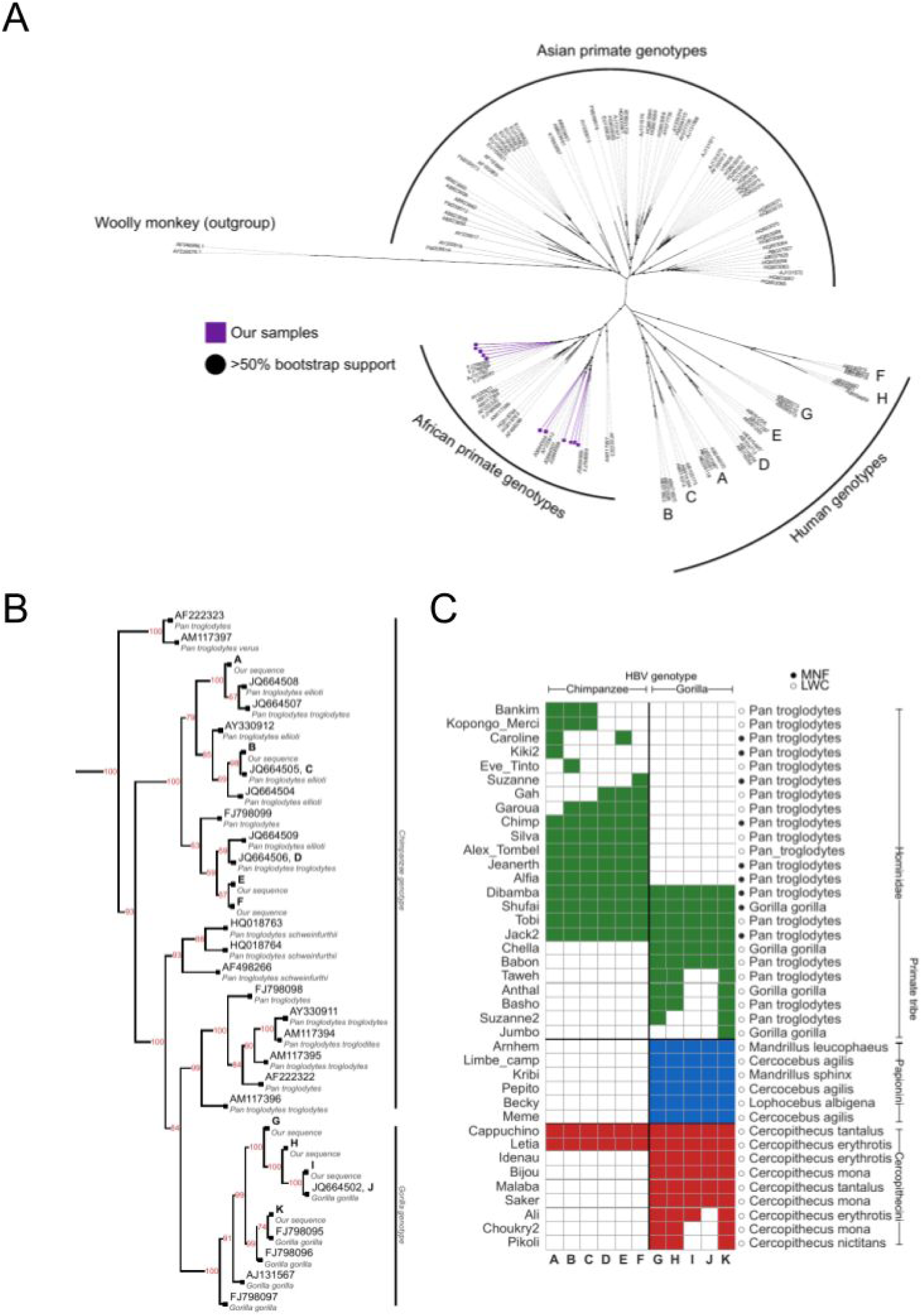
Hepatitis B virus in non-human primates. A) Unrooted phylogenetic tree of primate Hepatitis B virus sequences, with the three main clades and outgroup marked. Our sequences all fall within the African primate genotypes. B) Rooted tree of the African primate genotypes, with annotation for the split between chimpanzee and gorilla genotypes and bootstrap values on each branch. Leaves labeled A-K are our sequences. C) Heatmap showing the possible genotypes detected in each individual with reads aligning to HBV. Multiple entries in a row indicate that the genotype could not be disambiguated based on the sequence coverage.

**Supplementary figure 9:**
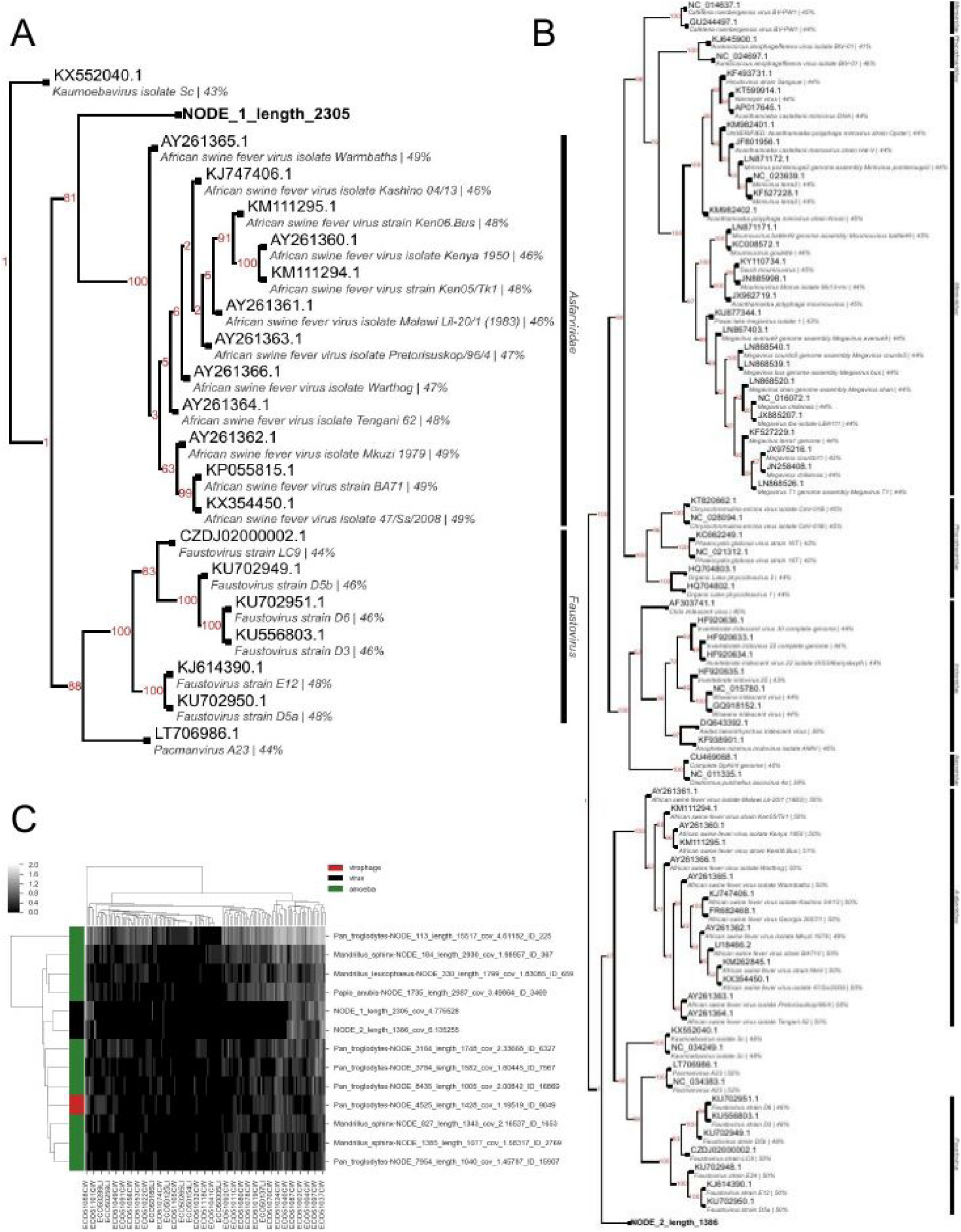
NCLDV trees and coverage. A) The gene from the longer of the two contigs is a cousin of African Swine Fever viruses, related to the Faustoviruses and Pacmanviruses. Numbers on branches are bootstrap values, percentage identities (compared to our sequence) are given after the name of the sequences. B) Tree made from the second of the genes, which resembles an RNA polymerase subunit. This sequence is a cousin of the Asfarviridae and Faustoviruses. C) Heatmap showing coverage (log10) of these contigs (black rows) along with amoeba sequences (green rows) and a virophage (red rows).

**Supplementary figure 10:**
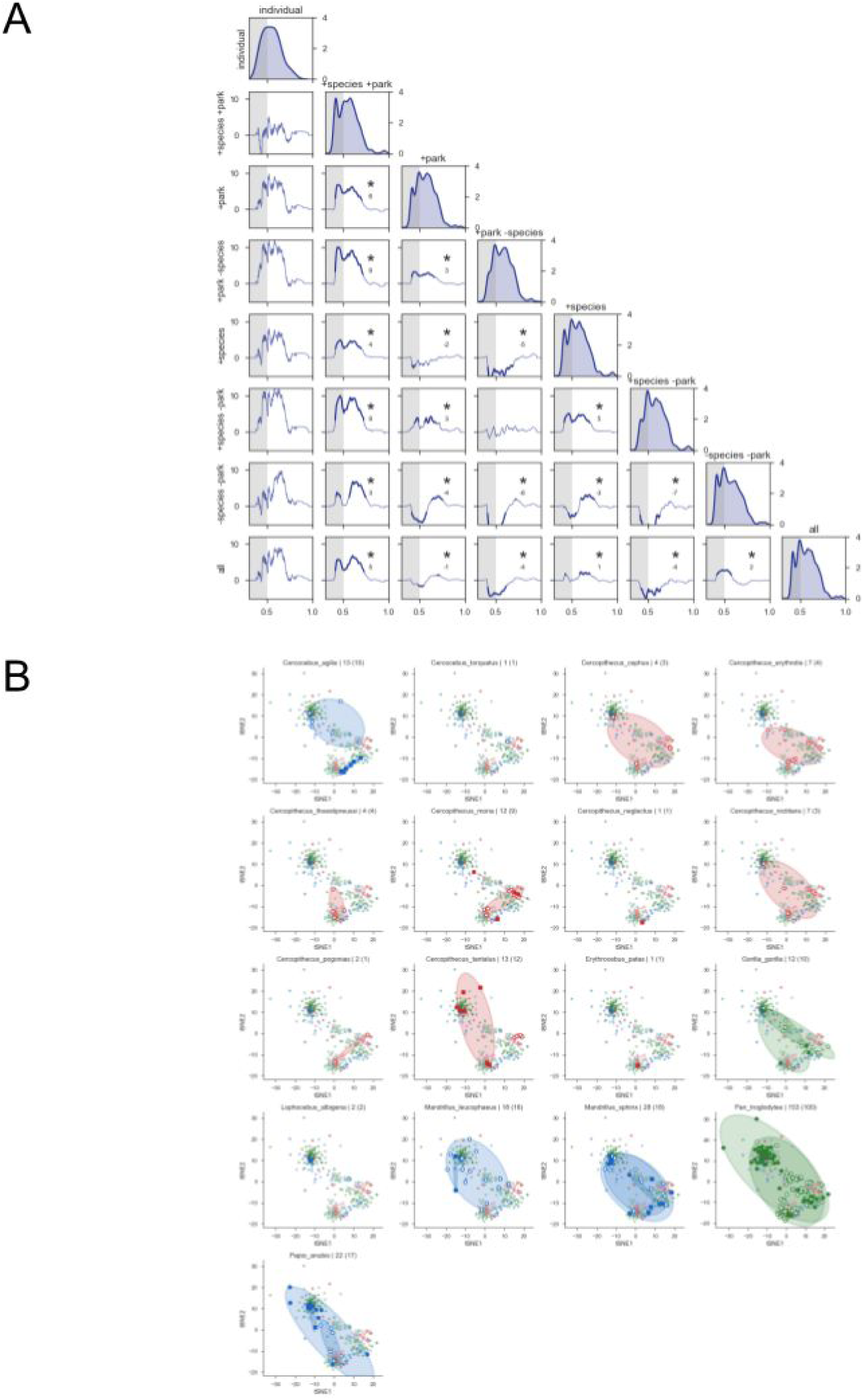
Multilevel UniFrac distance distributions and visualizations between sample subsets. A) Pairwise comparisons of KS statistics for different subsets of samples (lower triangle) together with the distance density (diagonal). Entries with “*” achieved statistical significance (p < 0.01) in the region highlighted (distance < 0.5, chosen as “low distance/high similarity”. Numbers indicate the magnitude of the maximum difference. B) UniFrac distances projected onto two dimensions using tSNE. Each graph has the samples from the species indicated highlighted, with ellipses containing the samples from each park. Open circles are samples from the Limbe Wildlife Centre, solid circles are from the Cameroon Wildlife Aid Fund. Colors are the tribe, as in figure 1A..

**Supplementary figure 11:**
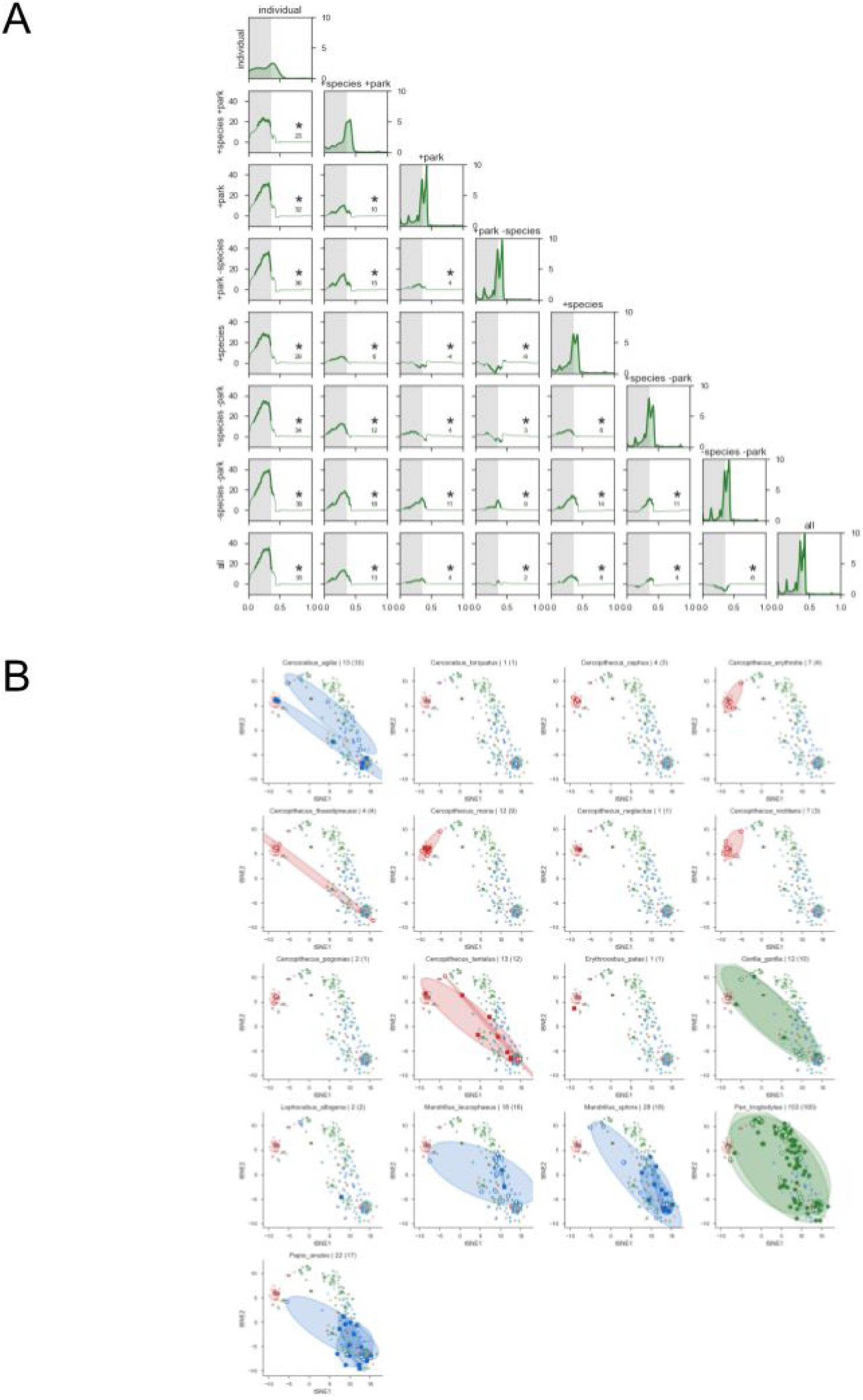
As figure S8, with taxa restricted to viruses.

